# How peak knee loads are affected by changing the mass of lower-limb body segments during walking

**DOI:** 10.1101/2025.01.29.635431

**Authors:** Delaney E. Miller, Ashley E. Brown, Nicholas A. Bianco, Rucha Bhise, Scott L. Delp, Steven H. Collins

**Affiliations:** Department of Mechanical Engineering, Stanford University, Stanford, California, United States of America; Department of Bioengineering, Stanford University, Stanford, California, United States of America; Department of Orthopaedic Surgery, Stanford University, Stanford, California, United States of America

## Abstract

For individuals with knee osteoarthritis, increased knee loading is linked to disease progression and pain. Some approaches to treating osteoarthritis, such as specialized footwear, braces, and powered exoskeletons, also increase the mass of the lower limbs, which could lead to increases in knee loads. Prior studies have investigated the effect of changes in torso mass and total body mass on peak knee contact forces, but the effects of increased leg mass remain unclear. In this study, we created musculoskeletal simulations informed by experimental data to estimate tibiofemoral knee contact force under different lower-limb segment mass conditions. The mass of the foot, shank, and thigh were varied by adding weights to each segment, separately and concurrently, as healthy young adults (N = 10) walked on a treadmill. Kinematics, kinetics, and muscle activity were recorded. Our simulations used an optimal control framework that enforced experimental kinematics while minimizing a combination of net joint moment errors and mismatch between measured and estimated muscle activity. The simulations revealed that adding mass to the lower-limb segments linearly increased early- and late-stance peaks in knee contact force, but that the slope of this relationship was different for each peak and each mass placement location. For each 1% of body weight (BW) added per limb (2% BW total) at the thigh, shank, and foot, early-stance peak knee contact force increased by 1.5%, 2.1%, and 5.9%, while late-stance peak contact force increased by 1.6%, 0.9% and 3.0%, respectively. Adding mass to the thigh and shank increases peak contact force at or below the rate of increase in body mass, while adding mass to the foot disproportionately increases peak knee contact force. These detrimental effects should be considered when designing interventions for osteoarthritis.

**Author summary:** For individuals with knee osteoarthritis, increased knee loading is linked to disease progression and pain. Some approaches to treating knee OA, such as specialized footwear, braces, and exoskeletons, increase the mass of the lower limbs, which could lead to increases in knee loads. However, the effects of changing lower limb segment mass on knee loading remain unclear. In this study, we created computer simulations informed by experimental data to estimate knee loads while walking with different amounts of added mass to the lower limb segments. We collected experimental data from 10 healthy participants while they walked on a treadmill with various amounts of mass added to their lower limb segments (thigh, shank, foot). We used these data and a biomechanically accurate model of each participant to construct realistic computer simulations, from which we obtained estimates of internal knee forces that contribute to total knee load. We found that peak knee load increased linearly with amount of added mass, but that this relationship differed by segment. Adding mass to the feet had the greatest effect on knee contact force. Our results can help to inform the design of interventions for knee osteoarthritis.

## Introduction

Knee osteoarthritis (OA), a degenerative condition of the knee joint, affects over 360 million adults worldwide and is becoming increasingly prevalent [1], underscoring the need for novel treatment approaches to slow disease progression. Several existing and emerging treatment approaches for knee OA aim to reduce joint loading, which is correlated with OA disease progression and pain [2–6]. For overweight or obese individuals with knee OA, reducing total knee load through weight loss can lead to clinically meaningful improvements in knee pain, function, and quality of life [7–9].

Interventions such as unloader braces, wedged insoles, mobility aids (e.g. canes or walkers), and gait modifications target knee load reduction or redistribution between the medial and lateral compartments of the knee joint [10–14]. However, these load-shifting approaches have limited effects on total knee load. Although relatively unexplored, a promising direction of knee OA treatment is the use of assistive exoskeletons to reduce knee loading [15–17]. It is possible that advancements in exoskeleton development could lead to larger reductions in knee joint load than existing approaches.

Understanding how worn mass affects peak knee contact force could inform the development of novel interventions for knee osteoarthritis such as lower-limb exoskeletons. Knee contact force results from a combination of intersegmental (or resultant) forces and compressive muscle forces from knee-spanning muscles. Muscle contributions to knee contact force cannot be measured directly, making it difficult to predict how adding mass to different locations on the body might affect knee loading during walking. Potential load-reducing treatment approaches such as exoskeletons, knee braces, and custom footwear require the user to wear some amount of mass on the legs, which introduces a tradeoff between the benefit of the intervention and the penalty of added mass. It is important to better understand how adding mass to the body affects peak knee contact force during walking, as these relationships can inform the design of assistive devices for individuals with knee OA.

Changes in total body weight and torso loading affect peak knee contact force. In the absence of kinematic changes, knee contact force is expected to change at a similar rate to changes in total body mass [18]. However, humans may modify their gait in response to changes in distribution or total magnitude of body mass. Aaboe et al. (2011) found that 13.5% weight loss corresponded to a 7% reduction in estimated peak knee contact force in a cohort of obese individuals with knee OA [9]. Weight loss may not map to an equivalent change in peak knee contact force because many obese adults adopt compensatory walking strategies, such as walking more upright than lean adults, to avoid increasing knee joint torque and power, and associated muscle forces [19]. In a similar manner, Lenton et al. (2018) found that adding approximately 18 and 36% bodyweight (BW) to the torso had little effect on lateral knee contact force in early stance, leading to a smaller increase in total contact force than expected (4.0 and 7.5%, respectively) [20]. In late-stance, however, peak total contact force increases at a rate more similar to the rate of added load (16 and 28%, respectively).

The distribution and magnitude of worn mass during walking affects gait kinematics, kinetics, and energetics. Increasing mass worn on the torso leads to a nearly linear increase in mechanical work and metabolic energy expenditure [21]. As mass is placed more distally along the lower limb, metabolic rate and muscle activity increase [22, 23]. Increasing the effective inertia of the leg is thought to increase effort associated with leg swing initiation and propagation. Despite these changes in energetics, several studies report relatively unchanged kinematics across different magnitudes and locations of added mass on the pelvis and lower limbs [23–25]. With high magnitudes of pelvis, thigh, and shank loading, Fang et al. (2022) observed significant changes in hip, knee, and ankle peak joint angles, although these changes were small in magnitude (approximately 1-3 degrees) [26]. From these findings, it is hard to estimate how peak knee contact force will change with added mass magnitude and placement.

Very few studies have investigated how the distribution of added mass affects knee loading, and the effect of adding mass to the lower limbs remains unclear. Although adding mass more distally is expected to lead to larger increases in muscle activity during swing, distribution of mass worn on the swing limb may affect the muscle activity of knee-spanning muscles on the stance limb, which could increase knee contact force. On the other hand, mass worn above the knee joint (e.g. on the thigh) on the stance leg is expected to contribute more to intersegmental loading than mass worn on the stance leg shank or foot [27]. The complex interaction between muscle and intersegmental contributions to knee loading make it difficult to predict how adding mass to the lower limbs will affect total knee joint contact force. Recordings from individuals with force-sensing knee implants have shown that wearing footwear increases early-stance knee load by 2-5% over walking barefoot, which suggests that adding mass to the foot may have a disproportionate effect on knee contact force [28]. However, specific footwear can simultaneously decrease late stance knee load by 6-9%, demonstrating the importance of considering the tradeoff between an intervention benefit and mass penalty.

It can be challenging to evaluate changes in knee loading because contact force cannot be measured in the intact knee. Although an instrumented knee prosthesis can be used to measure joint loads in vivo, these devices are extremely rare and only implanted in a select number of patients who undergo total knee arthroplasty. Instead, state-of-the-art methods use musculoskeletal models to estimate muscle forces and compute contact force [29–31]. Electromyography (EMG)-informed modeling approaches are an attractive approach to estimating knee contact force because they account for experimentally observed muscle coordination strategies [32–40]. These approaches usually solve for muscle excitations or forces that minimize a combination of control effort and EMG tracking error, subject to experimental kinematics and the parameters defined by a subject-specific anatomical model. The information encoded in EMG signals can be especially useful when individuals participate in a novel task and may not choose muscle activation patterns that solely minimize effort [41]. Although simulation-based estimates of knee contact force are subject to several sources of modeling and measurement error, some effects of error can be mitigated by evaluating intrasubject changes across conditions rather than comparing absolute values of contact force between subjects [42].

The purpose of this study was to evaluate the effects of adding mass to the thigh, shank, and foot on estimated peak knee joint contact force. Participants walked at a fixed speed on a treadmill while wearing different amounts of mass on their thighs, shanks, and feet. From experimental motion capture, force plate, and electromyography data, we generated EMG-informed musculoskeletal simulations using OpenSim Moco [39] to estimate total tibiofemoral knee contact force. We hypothesized that added mass would increase peak knee contact force, and that the increase in peak contact force would be greater for more distal load placement. Furthermore, we hypothesized that the effects of added mass on peak knee contact force would be both linear and additive; that is, increasing worn mass on a segment would proportionally increase peak contact force, and that the effect of adding mass to multiple lower limb segments would be similar to the summed effects of the individually worn masses. We expect these results to further our understanding of the relationship between mass distribution and knee loading, which will inform the development of assistive devices for knee osteoarthritis.

## Materials and methods

### Participants

Ten healthy adults (4 M, 6 F; 1.71 m, 65.5 kg, 27 y.o.) participated in this study. The study protocol was approved by the Stanford University Institutional Review Board, and all participants provided written informed consent before participating in the study. Individuals were excluded if they had any ongoing orthopedic conditions, such as arthritis or ligament injuries to the knee.

### Data collection

Participants performed 11 different trials during a single data collection session: a calibration trial, an initial unloaded walking trial, 8 loaded walking trials, and a final unloaded walking trial. Surface electromyography (EMG) signals (Delsys Inc., Boston, MA USA) were collected at 1250 Hz from 9 lower-limb muscles on the right leg: soleus (medial aspect), medial gastrocnemius, lateral gastrocnemius, tibialis anterior, semitendinosus, biceps femoris long head, vastus medialis, vastus lateralis, and rectus femoris. Before placing each electrode, an experimenter shaved and cleaned the electrode location with isopropyl alcohol. After electrode placement, participants performed a series of maximum voluntary contraction (MVC) exercises that were later used to normalize EMG data.

Thirty-two retroreflective markers were placed on anatomical landmarks on the arms, torso, pelvis, thighs, lower legs, and feet. Additionally, 24 markers were used to aid in limb tracking. Six markers that were placed on anatomical landmarks were removed after the calibration trial due to potential interference during walking. Marker data were collected using a 12-camera optical motion capture system at 100 Hz (Vicon Motion Systems Ltd, Oxford UK). Ground reaction forces collected from a split-belt treadmill (Bertec Corporation, Columbus, OH, USA) at 1250 Hz.

Participants first performed a static and range-of-motion calibration trial, which we used to estimate hip joint centers and tracking marker reference locations. Participants then completed a series of 10 walking trials in which they walked for 5 minutes at a speed of 1.25 m/s on a split-belt treadmill (Fig 1). We recorded data from the last 80 seconds of each gait trial and averaged across stance phases to obtain our outcome measures for each condition. In the first and final walking trial, participants walked without any additional mass. In a random order, participants completed sets of thigh, shank, and foot loading trials. Each set consisted of two trials in increasing mass order. Mass was added as a function of participant body weight (BW). We added 2% and 4% of BW to each thigh, 1.5% and 3% of BW to each shank, and 1% and 2% of BW to each foot. These magnitudes were chosen to span the range of masses of current and emerging assistive devices intended to address knee loading. Finally, participants completed two loading trials in which mass was placed on all three segments using the lower and higher magnitudes for each segment (4.5% and 9% BW per leg). For a 70 kg person, the added mass ranged from 0.7 kg (1% BW at the foot in the low condition) to 6.3 kg (9% BW across all segments in the high condition) per leg.

**Fig 1.**
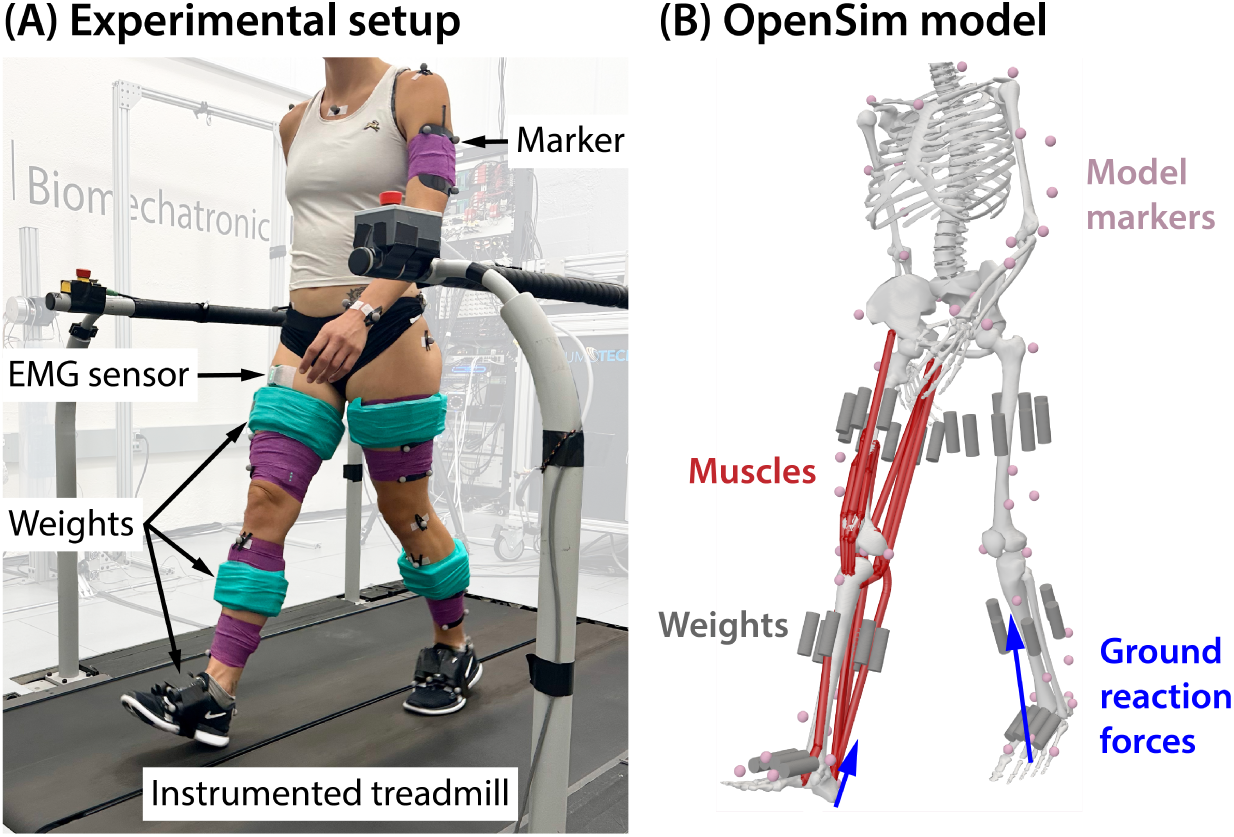
Experimental setup. A. Participants walked on an instrumented treadmill while wearing varying amounts of mass on their thighs, shanks, and feet. Electromyography (EMG), marker data, and ground reaction forces were collected. B. Experimental data were used to construct musculoskeletal simulations in OpenSim [43].

Mass was added to each lower-limb segment using 3-inch-long stainless-steel rods of 1-inch diameter (300 g) and 0.5-inch diameter (75 g). For the thigh and shank, rods were aligned with the limb segment and distributed evenly around the limb circumference, except on the medial thigh to avoid interference during walking. Rods were placed at the approximate mass center along the proximal-distal axis of the limb segment [44]. For the foot, rods were placed on the dorsal surface of the midfoot and aligned with the metatarsals. Masses were adhered to the participant using a combination of hook-and-loop closures and duct tape. Rod combinations were selected to most closely match the desired percentage of body mass, which resulted in a maximum discrepancy of approximately 37.5 grams. For our smallest participant in the lowest loading condition, this discrepancy would amount to approximately 7% error.

### Data processing

Experimental data were filtered and segmented into individual stance phases of the right leg. Electromyography signals were high-pass filtered at 20 Hz, full wave rectified, then low-pass filtered at 6 Hz. A 250-millisecond sliding window average was used to compute maximum signals from MVC trials. EMG data from gait trials were normalized by the maximum value seen across MVC and gait trials for that muscle such that no signals exceeded 1. EMG signals were shifted by a delay of 53 milliseconds to account for the electromechanical delay between neural excitation and muscle force production [45]. Experimental recordings were visually inspected to remove erroneous data resulting from motion artifact or sensor attachment issues. Ground reaction force data were low-pass filtered at 8 Hz.

We estimated joint kinematics and kinetics using OpenSim 4.4 [43]. We used a generic full-body musculoskeletal model [46] with updated muscle paths [14] that had 35 degrees of freedom and 80 lower-extremity muscles. Torque actuators drove the torso and upper extremities to achieve dynamic consistency. For our EMG-informed simulations, we removed all but 12 musculotendon units on the EMG-instrumented (right) lower limb. In addition to the nine muscles for which we collected surface EMG data, we mapped three signals to adjacent muscles [33, 47]: vastus medialis to vastus intermedius, biceps femoris long head to biceps femoris short head, and semitendinosus to the semimembranosus. The remaining muscles were replaced with ideal torque actuators for each degree of freedom.

The modified generic model was scaled using marker data from the static pose in the calibration trial. Hip joint centers were estimated from the range-of-motion tasks during the calibration trial to additionally inform pelvis scaling. We performed an optimization procedure to preserve the operating ranges for each muscle-tendon actuator as described in [48, 49]. We scaled peak isometric muscle forces to the mass and height of each participant using the relationships established by Handsfield et al. [50]. The steel rods used in each weighted walking condition were modeled in OpenSim and attached to the appropriate segment (Fig 1b). We calculated joint kinematics from marker data using OpenSim’s Inverse Kinematics tool. Average root-mean-square marker error (*<*3 cm) and maximum marker error (*<*5 cm) fell within recommended guidelines [51]. We filtered kinematic data at 8 Hz before computing inverse dynamics to obtain net joint moments.

### EMG-informed simulation

We created EMG-informed simulations in OpenSim Moco to estimate muscle forces, which were then used to compute knee contact force (Fig 2). Experimental kinematics and ground reaction forces were prescribed. We used a two-stage simulation approach in which we first calibrated the magnitudes of experimental EMG signals (“Calibration”), then used the calibrated signals to process the remainder of the trials (“Execution”). For each simulation, we solved for the muscle excitations and reserve torques that minimized muscle excitation effort and reserve torque effort while tracking experimental EMG. Reserve torques at the right knee and ankle were heavily penalized, while effort across the remaining reserve torques was assigned a small weight to encourage optimizer convergence. The cost function can be represented by the following equation:

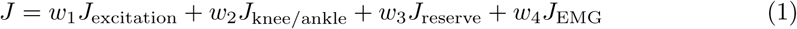

**Fig 2.**
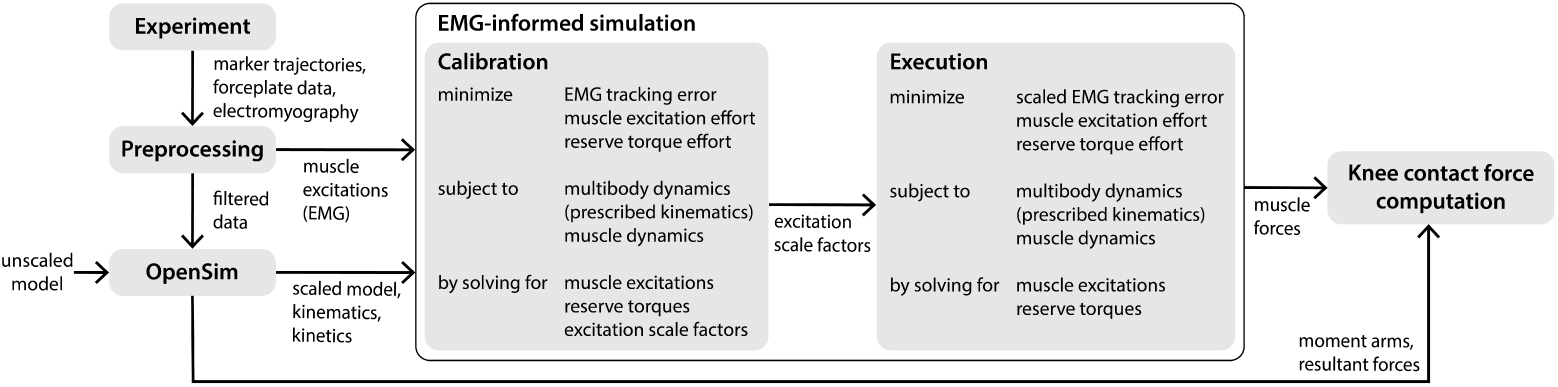
Simulation pipeline. Experimental data were preprocessed and passed into OpenSim to scale an anatomical model and generate kinematic and kinetic data. Muscle excitations from experimental EMG data were passed directly to the EMG-informed simulations in OpenSim Moco. We employed a two-stage EMG-informed simulation approach in which we prescribed experimental kinematics and dynamics. In both stages, we solved for adjusted muscle excitations and reserve torques that minimized a weighted sum of EMG tracking error (sum of squared errors between EMG and simulation-adjusted excitations), muscle excitation effort (sum of squared muscle excitations), and reserve torque effort (sum of squared reserve torques). Reserve torques at each joint provided additional strength in the model to enforce dynamic consistency; minimizing reserve torque effort was conceptually equivalent to minimizing net joint moment error. During Calibration, a subset of walking data (split into individual stance phases) was used to calibrate the magnitude of muscle excitations from experimental EMG such that muscle-generated moments most closely matched the observed net joint moments. During Execution, experimental excitations were adjusted using scale factors obtained during the Calibration stage. Muscle forces were computed from simulation-adjusted muscle excitations and the model state. Knee contact force was computed from a combination of muscle and resultant (intersegmental) forces.

Each term was integrated over a single stance phase of the right leg with initial time *t*_0_ and final time *t_f_*. Muscle excitation effort, *J*_excitation_, is the sum of squared muscle excitations, integrated over stance. Reserve torque effort for the right knee and ankle, *J*_knee*/*ankle_, is the sum of squared knee and ankle reserve torques, integrated over stance. Similarly, the reserve torque effort at all other joints, *J*_reserve_, is the sum of squared reserve torques for all other joints in the model, integrated over stance. Finally, the experimental EMG tracking error, *J*_knee*/*ankle_, is the sum of squared error between each time-delayed experimental muscle EMG signal and the adjusted muscle excitation found by the optimizer. The weights *w* on these terms varied between Calibration and Execution, as described below.

### Calibration

To better match muscle-driven moments with the net joint moments obtained through inverse dynamics, we employed a calibration step to further scale the experimental EMG data. EMG data were previously normalized to the maximum value seen across all maximum voluntary contraction (MVC) and gait trials. However, MVC efforts are commonly submaximal, which results in normalized EMG signals that are too large in magnitude. In the simulation setup, an additional scale factor parameter was introduced for each muscle such that each reference EMG signal could be scaled by a factor between 0.5 and 1. We solved for adjusted muscle excitations, reserve torques, and EMG scale factors that minimized the objective function described in Eqn 1. For each participant, we ran simulations on 10 strides from the unweighted walking condition and obtained a set of scale factors from each stride. These scale factors were averaged across the 10 strides to produce a single scale factor for each muscle, which was used to scale the reference EMG data during the Execution stage.

During Calibration, a high weight *w*_2_ was assigned to knee and ankle reserve torque effort such that the muscles in the model were primarily responsible for producing the observed joint moments. A high weight *w*_1_ was also assigned to excitation effort to prevent submaximal MVC signals from artificially inflating the experimental EMG signals. Because we expected minimal co-contraction during normal walking in a healthy population, this penalty scaled down the reference EMG signals if needed. In both Calibration and Execution, reserve torque effort at the remaining joints (*J*_reserve_) was assigned a small weight to encourage optimizer convergence but did not contribute significantly to the objective function value.

### Execution

For execution, reference EMG signals were adjusted by the scale factors determined in the calibration step. The weight on knee and ankle reserve actuator effort *w*_2_ was decreased to allow for slightly more deviation between the muscle generated joint moments and results from inverse dynamics, especially near heel strike and toe off where center-of-pressure calculations have reduced accuracy. The weight on excitation effort was decreased to a nominal value, as the scale factors obtained during Calibration determined the overall magnitude of the solution excitations. The weight on EMG tracking error *w*_4_ was held constant across Calibration and Execution. However, adjusting other weights in the cost function increased the relative importance of minimizing EMG tracking error, such that variation in experimental EMG signals was more accurately captured.

Each simulation produced a solution with adjusted muscle excitations and reserve actuator torques. The muscle excitations, along with known musculotendon states from the scaled anatomical model and prescribed kinematics, were used to compute muscle forces.

### Reserve torque actuators

Reserve torque actuators at the right knee and ankle provided additional strength in the model to enforce dynamic consistency despite errors arising from the data. Penalizing reserve torque effort in the optimization cost function was conceptually equivalent to minimizing muscle-driven joint moment errors. At the knee and ankle, reserve torque effort was heavily penalized in the cost function such that the muscles in the model were primarily responsible for producing the joint moments arising from inverse dynamics. For degrees of freedom with no muscle actuation, torque actuators supplied the full joint moment needed to enforce the dynamic constraints for motion. Torque actuators at these degrees of freedom, along with degrees of freedom that were partially actuated by the muscles (e.g. hip flexion), were assigned a small effort weight to encourage optimization convergence.

### Computation of total knee contact force

We computed knee contact force using the JointReaction analysis in OpenSim, as described in detail by Steele et al. (2012) [30]. This tool computes the forces and moments carried by the joint structure as a result of all structures and forces that produce the desired joint kinematics (muscles, cartilage, ligaments, reserve actuators). The muscle forces estimated from the EMG-informed simulations, along with muscle moment arms and resultant (intersegmental) forces were used to compute compressive tibiofemoral contact force. Total knee contact force was taken as the axial component in the tibia reference frame. Early- and late-stance peaks in knee contact force were taken as the maximum values within 15-40% of stance and 60-90% of stance, respectively.

We also computed the intersegmental (or resultant) contribution to total knee contact force for each subject. Equivalently, this analysis allowed us to extract the contribution of tensile muscle forces to knee joint loading. For each condition, we replaced all of the muscles in the model with ideal torque reserves at each joint. These torque reserves supplied the necessary joint moments to produce the observed motion without contributing to axial knee joint loading. We then used the JointReaction analysis tool in OpenSim to compute the intersegmental contribution to tibiofemoral contact force.

### Statistical analysis

The sample size of this study was informed by an a priori power analysis. We would expect under the most basic assumption that increasing body mass by 1% would increase peak knee contact force by the same percentage. Because the minimum amount of added mass was 2% BW (1% per leg, worn at the foot), we were interested in detecting a minimum change of 2% in peak contact force. We assumed a standard deviation across participants of 2% BW. A sample size of 10 participants yielded a statistical power of 0.8 (two-tailed t-test, alpha = 0.05).

Because our primary outcome was a linear regression model relating segment mass to change in peak knee contact force, we conducted a power analysis for a linear model with three predictors (mass at thigh, mass at shank, and mass at foot). We assumed a sample size of 60 observations based on testing 6 conditions with 10 participants.

Because we computed a percent change in peak contact force relative to the unweighted condition, we assumed all observations were independent. For a statistical power of 0.8 with a significance level of 0.05, the minimum proportion of variance explained by the model (R-squared) would be 0.17. Thus, we could detect a small effect of added mass on percent change in peak knee contact force.

To evaluate how mass at each segment affected the measured outcomes, we fit linear mixed-effects models to the data (fixed effects: mass at thigh, mass at shank, mass at foot; random effect: subject) to account for individual subject differences. In addition, we fit a simple linear regression model (*y* = *a* ∗ *x*) to describe the relationship between added segment mass (as a percentage of body weight) and percent change in peak knee contact force; models were fit separately for the early- and late-stance peaks. Measured outcomes were taken as the average across stance from the final 80 seconds of each 5 minute walking condition. The significance level for all analyses was alpha = 0.05. Statistical analysis was performed using MATLAB (Mathworks, Inc.).

## Results

### Simulation results

During the region of interest (10-90% of stance), reserve torque actuators at the knee and ankle supplied negligible amounts of torque, such that the net joint moments were driven almost entirely by muscles. The knee and ankle reserve actuators supplied average peak torques of 0.028 Nm/kg and 9e-3 Nm/kg, respectively, normalized to participant body mass. These values corresponded to 5% of the peak knee joint moment and 0.5% of the peak ankle joint moment in the region of interest.

The adjusted muscle excitations closely tracked the experimental EMG signals. In the region of interest (10-90% of stance), the average root-mean-square error between experimental and adjusted excitations was 0.02, where excitations could range from 0 to 1. Additional simulation results are reported in S1 Appendix.

### Peak knee contact force

When controlling for subject, magnitude of added mass at each segment had an effect on peak knee contact force, both for the early-stance peak (linear mixed-effects model, p *<* 0.001) and the late-stance peak (linear mixed-effects model, p ≤ 0.001). Across segments, adding mass increased peak knee contact force for both peaks (Fig 3). Data were included from all walking conditions, including the condition with no added mass. Subject was included as a random effect to account for individual subject offsets. Peak knee contact force and added mass at each segment were normalized to participant body mass and expressed in body weight (BW).

**Fig 3.**
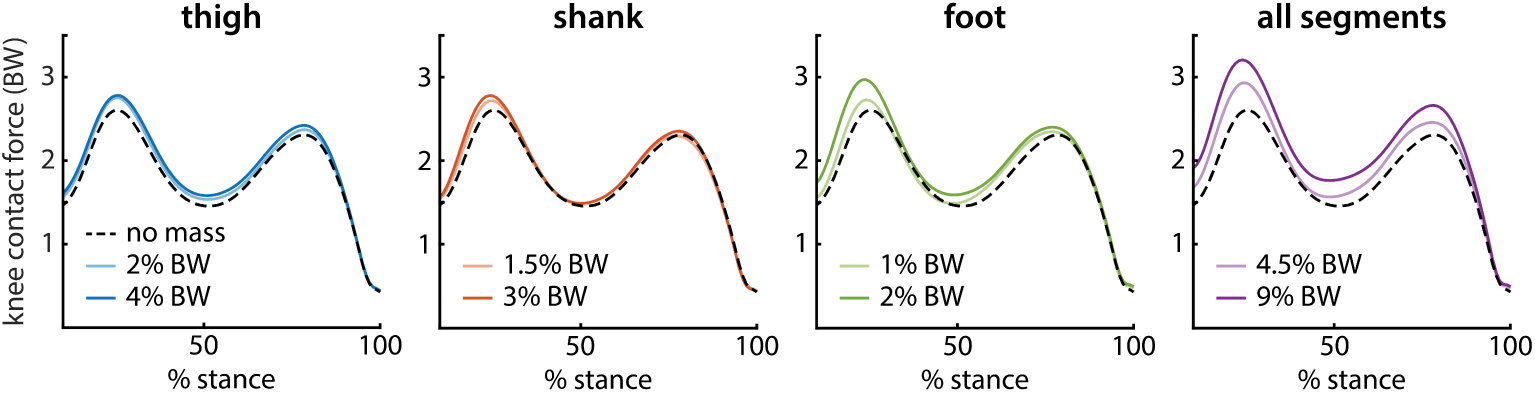
Knee contact force across loading conditions. Total knee contact force is shown as a percentage of stance for all loading conditions. On each plot, the unloaded (no mass) condition is represented as a dashed black line. The light and dark colored lines represent the low and high mass loading conditions, respectively, for each segment.

The relationship between added mass at each segment and percent change in peak knee contact force is well approximated by a line (Fig 4). To normalize data across participants, we computed percent change in peak contact force from the unweighted (no mass) condition for each subject. We fit a simple linear regression model (*y* = *a* ∗ *x*) to percent change in peak contact force for both early- and late-stance peak (Table 1). Added mass per leg was expressed in percent body weight (% BW). Data from the no mass condition and all segment loading conditions were included in these models. For the early-stance peak in knee contact force, the model is given by:

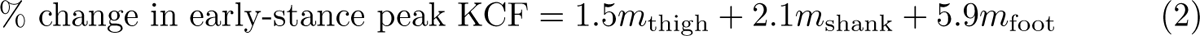

**Fig 4.**
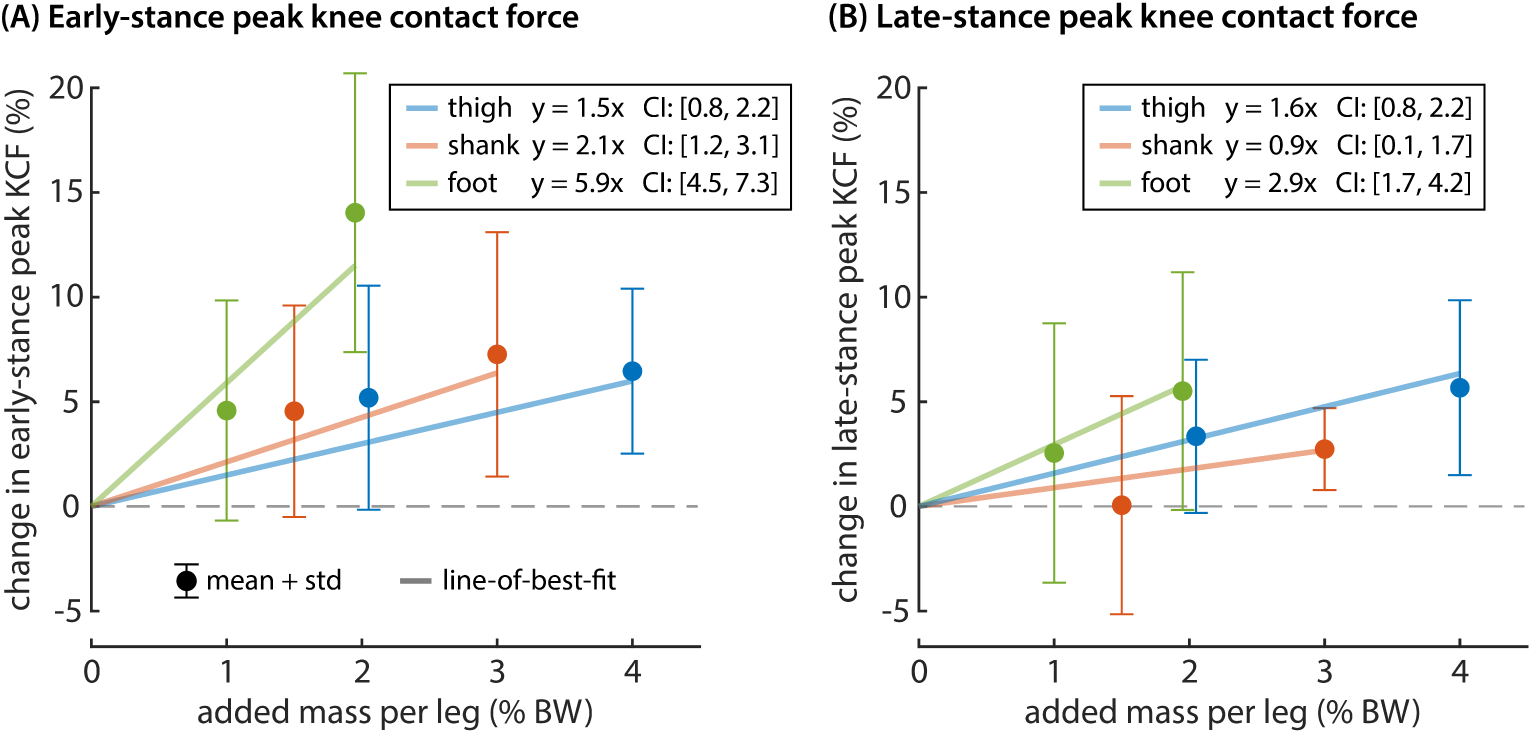
Relationship between added mass and changes in peak knee contact force. Percent change in (A) early- and (B) late-stance peak knee contact force was computed relative to the unloaded condition. The mean and standard deviation across participants is shown for each of the single-segment loading conditions. We fit a simple linear model to added thigh mass, shank mass, and foot mass using data from all eight loading conditions. The portion of this model associated with each segment is plotted as a line-of-best fit over the data (p *<* 0.032).

**Table 1.** Summary of two linear models relating added mass at each segment (expressed as % BW per leg) to percent change (%Δ) in early- and late-stance peak knee contact force (KCF), separately. The coefficient value, 95% confidence interval, and p-value is presented for each added mass segment.

| added mass segment | %$\Delta$ in early-stance peak KCF<br>(Eqn 2) | | | %$\Delta$ in late-stance peak KCF<br>(Eqn 3) | | |
| --- | --- | --- | --- | --- | --- | --- |
|  | coefficient value | confidence interval | p-value | coefficient value | confidence interval | p-value |
| thigh | 1.5 | [0.2, 2.2] | <0.001 | 1.6 | [1.0, 2.2] | <0.001 |
| shank | 2.1 | [1.2, 3.1] | <0.001 | 0.9 | [0.1, 1.7] | 0.032 |
| foot | 5.9 | [4.5, 7.3] | <0.001 | 3.0 | [1.7, 4.2] | <0.001 |

Each term had a strong association with change in early-stance peak knee contact force (p *<* 0.001). For the late-stance peak in knee contact force, the model is given by:

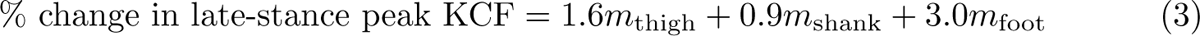

Each term had a strong association with change in late-stance peak knee contact force (p *<* 0.032).

Across loading conditions, muscle force contributions were primarily responsible for changes in peak knee contact force, rather than intersegmental force contributions (Fig 5). In early-stance, muscle contributions comprised between 65.3% (foot loading, low mass) and 99.6% (shank loading, high mass) of the observed change in peak knee contact force. In late stance, muscle contributions accounted for 57.4% (thigh loading, high mass) to 83.0% (foot loading, high mass) of changes in peak load.

**Fig 5.**
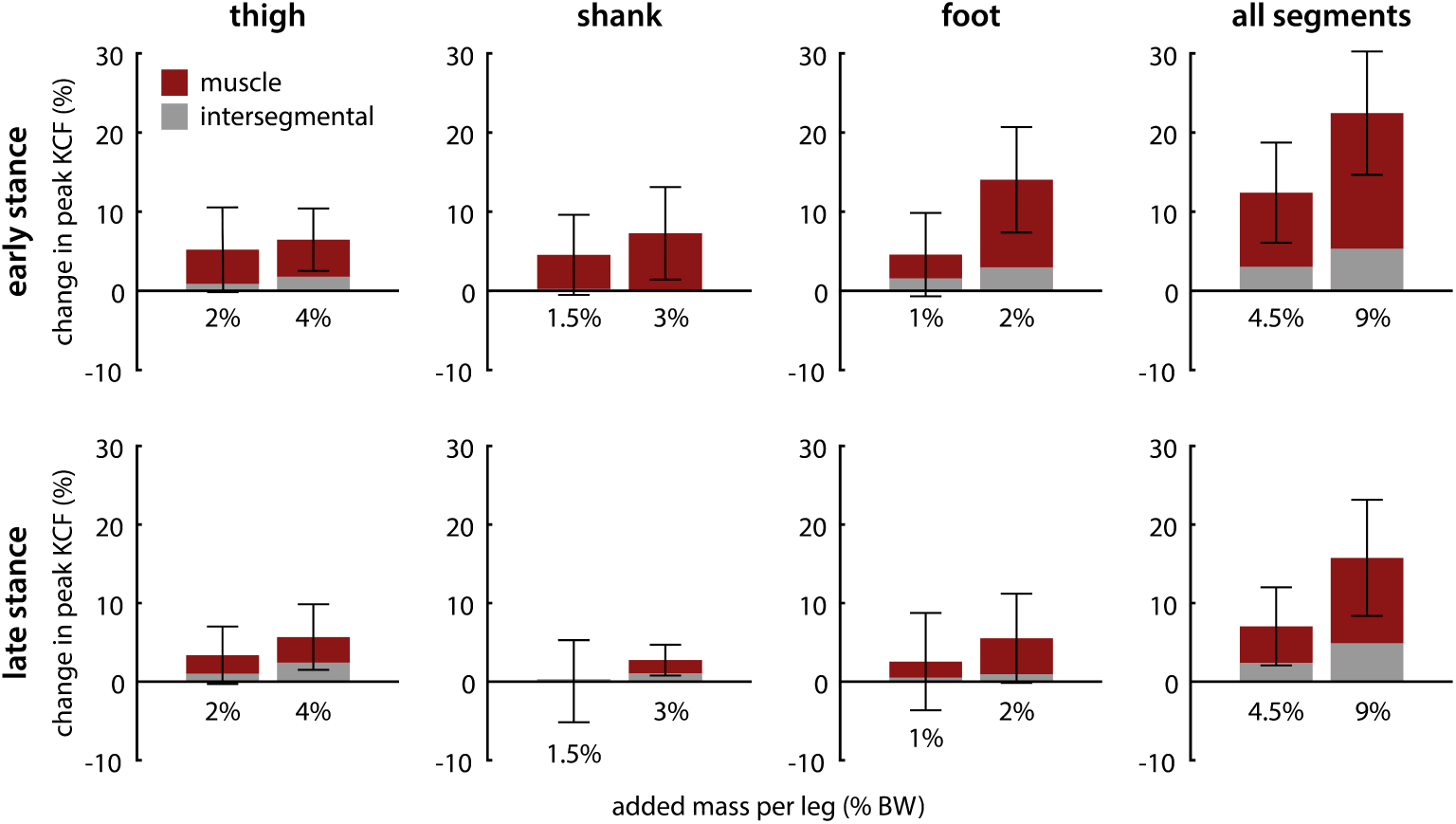
Muscle versus intersegmental contributions to changes in peak knee contact force. Muscle force contributions (shown in red) dominated total change in peak contact force across loading conditions. Each plot column represents a different segment loading condition. Standard deviation bars are shown across participants. *Top row:* Changes in early-stance peak contact force. *Bottom row:* Changes in late-stance peak contact force.

We trained a separate linear regression model on the individual segment loading conditions to evaluate the performance of this model on the unseen all-segment loading conditions. This test allowed us to assess whether or not there was an interaction effect when adding mass to multiple segments. We used the 95% confidence intervals from the model trained on individual segment loading conditions to estimate the upper and lower bounds we would expect on the group average outcome variables for the all-segment loading conditions. We found that for both low (4.5% BW per leg) and high (9% BW per leg) all-segment loading conditions, the estimated group average changes in peak contact force fell within the 95% confidence interval bounds, except for the the early-stance peak in the high loading condition (9% BW per leg). Our model trained on individual segment loading data overpredicted the change in early-stance peak contact force for this condition. The inter-subject average increase in contact force peak was 22.4%, while our model predicted a change of 27.9%. The lower end of the 95% confidence interval range predicted a minimum change of 22.6%. This result indicates that there may be an interaction effect between segment masses in early stance, such that the combined effect of added mass is smaller than the sum of the effects of the individual segment masses.

### Temporal, kinematic, and kinetic changes

We observed changes in temporal gait characteristics as well as knee kinematics and kinetics under different loading conditions (Fig 6). Adding mass to the foot and the thigh increased stride time (linear mixed-effects model, p *<* 0.001). We fit a simple linear regression model to estimate the effects of adding segment mass on percent change in stride time. We found that adding 1% BW per foot increased stride time by 2.5% (p *<* 0.001) and adding 1% BW per thigh increased stride time by 0.26% (p = 0.002), but adding mass to the shank did not (p = 0.49). Duty factor, the ratio of stance time to stride time, decreased with foot mass but slightly increased with thigh mass (linear mixed-effects model, p *<* 0.01).

**Fig 6.**
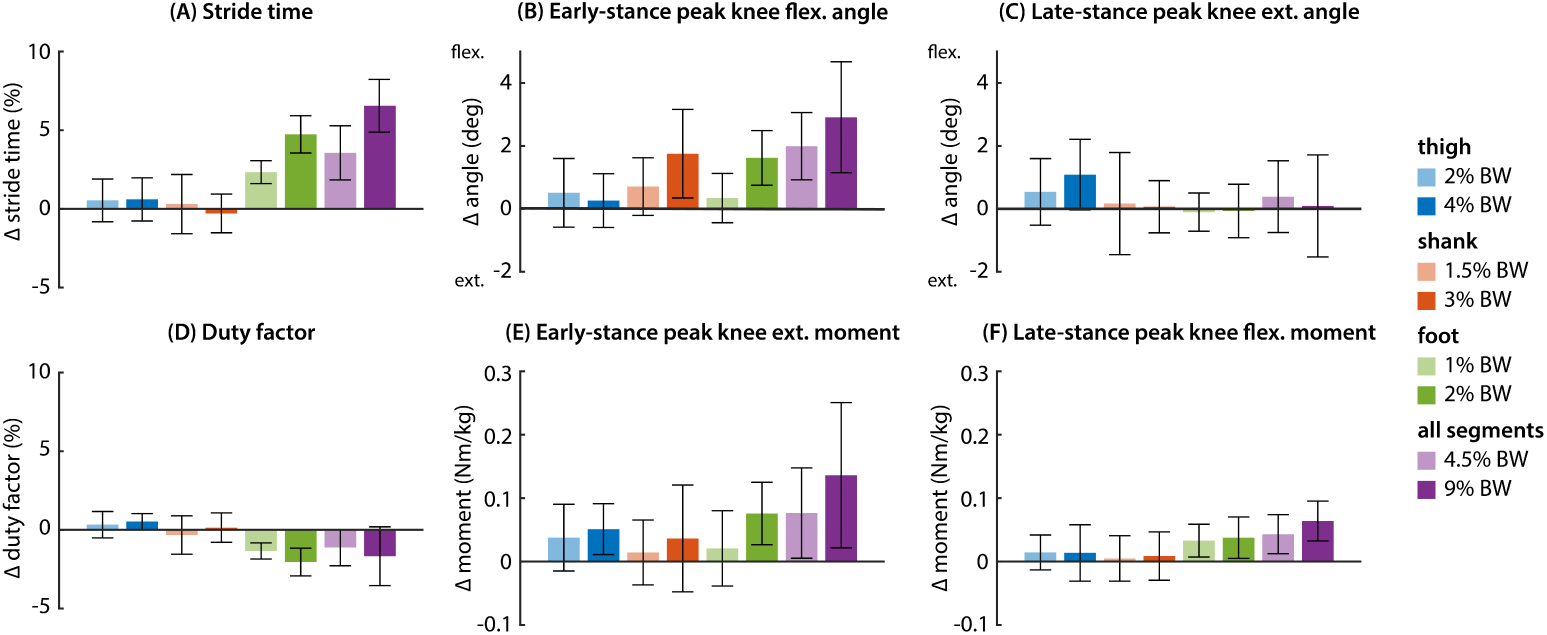
Changes in temporal gait characteristics, kinematics, and kinetics. All changes are shown relative to the unloaded (no mass) walking conditions. (A) Stride time increased with foot loading and, to a lesser extent, thigh loading. (B) Early-stance peak knee angle became more flexed with shank and foot loading. (C) Late-stance peak knee extension angle became more flexed with thigh loading. (D) Late-stance duty factor decreased with foot loading and increased slightly with thigh loading. (E) Magnitude of early-stance peak knee extension moment increased with thigh and foot loading. (F) Magnitude of late-stance peak knee flexion moment increased with thigh and foot loading. Changes in kinetics with thigh loading were explained by changes in total mass.

Some kinematic adaptations at the knee were observed across loading conditions. In early stance, peak knee flexion angle increased with shank and foot loading (linear mixed-effects model, p *<* 0.001). We found that increased knee flexion angle with foot loading was explained by an increase in stride time, and that this effect was no longer significant when data were normalized to stride time. In late stance, thigh loading decreased late-stance peak extension angle such that the knee remained more flexed (linear mixed-effects model, p = 0.004).

Kinetic changes at the knee were largely explained by increases in total worn mass and changes in stride time. Peak early-stance knee extension moment increased with thigh and foot loading (linear mixed-effects model, p *<* 0.001). Late-stance peak knee flexion moment increased with both thigh and foot loading (linear mixed-effects model, p *<* 0.02). We found that thigh loading no longer had an effect on peak joint moments when the outcome measures were normalized to total worn mass. Increased early-stance peak extension moment with foot loading was explained by a combination of total worn mass and increased stride time. Foot loading still increased late-stance peak knee flexion moment, despite normalizing to total worn mass and stride time (linear mixed-effects model, p *<* 0.001).

## Discussion

In this study, we found that magnitude of added mass to the thigh, shank, and foot had an effect on peak contact force. Regardless of mass placement location, the relationship between magnitude of added mass and percent change in peak contact force was well approximated by a line. The slope of this line differed across segments (thigh, shank, foot) and between early- and late-stance peak. The effects of adding mass to multiple lower-limb segments appeared to be mostly additive. However, there may be an interaction effect between high magnitudes of segment mass in early-stance, as all-segment loading led to a smaller increase in knee contact force than predicted by summing the single-segment contributions.

Adding mass to the feet had the greatest effect on both early- and late-stance peak knee contact force, with a larger effect on the early-stance peak. In early-stance, for each 1% BW added per foot, peak knee contact force increased by approximately 6%. In late stance, each 1% BW added per foot led to approximately a 3% increase. We found that foot mass increased stride time, which has been observed previously [23, 52]. In our study, stride time increased by approximately 2.5% for each 1% BW added per foot. We observed longer swing times with added foot mass, which can be attributed to reducing the natural frequency of the swing leg by increasing the effective inertia about the hip joint. At a fixed speed, increasing stride time, or, equivalently, stride length, has been shown to increase peak knee contact force at a similar rate [53, 54]. However, increased stride time only partially explains the increases in peak knee contact force we observed with foot loading, especially in early-stance.

Wearing mass on the contralateral (swing) foot may disproportionately increase peak knee contact force by increasing the inertial effects of the swing leg. Mass worn on the stance foot should have negligible effect on peak contact force, as the mass is worn below the knee joint and the foot moves very little at those timepoints in the gait cycle. However, in early-stance when the first peak of knee contact force occurs, the swing leg has recently pushed off the ground and trails the stance leg. The acceleration of the swing foot is relatively high at this point in the gait cycle because the foot needs to move farther forward during swing than any other part of the body. The forces to produce this acceleration must pass through both legs to the ground, requiring increased activation of stance-leg knee musculature and thereby increasing knee contact forces. These dynamics of the swing leg have previously been described [55] and used to explain why adding mass to the feet also leads to a disproportionate increase in the metabolic cost of walking [23].

Thigh mass had a moderate effect on both early- and late-stance peak knee contact force. For each 1% BW added per thigh, peak contact force increased by approximately 1.5%. Reasoning from basic principles, an increase of 2% BW (1% per leg) would be expected to lead to the same rate of 2% increase in peak knee contact force. Although we observed a smaller increase in peak contact force with thigh loading, a 2% increase still fell within the 95% confidence interval for our model. Muscle contributions to peak contact force made up the majority of the observed increases in peak contact force with thigh loading, rather than by intersegmental contributions (Fig 5). These results are consistent with previous findings that muscle contributions make up approximately 50-75% of total knee contact force during walking [29, 38].

Shank loading had a greater effect on early-stance peak contact force than late-stance peak contact force. In early-stance, each 1% BW added per shank increase peak contact force by approximately 2%. In late-stance, shank loading increased peak contact force at a rate of approximately 1% per 1% BW added per leg. Muscle contributions to knee contact force were almost entirely responsible for increases in peak contact force due to shank loading (Fige 5). Although placed more distally than thigh mass, shank mass may have had less effect on late-stance peak contact force due to its placement below the knee joint. On the stance leg, mass worn above the knee joint contributes directly to axial intersegmental load, while mass worn on the shank only contributes to intersegmental load through rotational inertia effects, which are small. Because the shank does not accelerate as much as the foot during swing, adding mass to the swing shank requires less activation of the stance leg muscles and, as a result, has less effect on knee contact force.

While adding mass to multiple segments appeared to have a mostly additive effect on peak contact force, there may be an interaction effect between segment masses in early stance. Our model trained on single-segment loading conditions predicted an increase in early-stance peak contact force of 28% with high magnitude all-segment loading (9% BW per leg), but the group average increase in peak contact force for this condition was only 22%. This result fell just outside the 95% confidence interval range for our model. While more data is required to support this conclusion, it is possible that adding mass to multiple lower-limb segments may be less costly than expected. We added higher magnitudes of mass more proximally, so the center of mass of the leg was less altered in the all-segment loading conditions than in the single segment conditions. A more proximal center of mass location than in the shank and foot loading conditions may have led to lower than expected inertial effects of the contralateral swing leg. There may have also been a motor learning effect in which the all-segment loading distribution felt more natural than the single segment loading distributions, despite the higher added mass.

This experiment had several limitations. We did not evaluate the effects of unilateral loading, which may be applicable to certain treatments or interventions, such as a unilateral leg brace or exoskeleton. Furthermore, participants were healthy, young adults, and it remains unclear how these findings might extend to an older population with knee OA, who may have altered gait characteristics [56–60]. However, estimating knee contact force using EMG-informed methods can be more challenging in this population, as obesity and muscle atrophy are frequently observed among individuals with knee OA [61–63], which can reduce the quality of collected EMG.

Due to the number of loading conditions tested, participants walked in each condition for five minutes, which may not have been enough time to adapt to each loading condition. During the high foot mass and high all segment loading conditions, we observed a small increase in early-stance hamstrings muscle activity, which may indicate that participants hadn’t fully adapted to the novel coordination task [41].

However, this increase in activation could also be associated with increased stride times. To reduce the effects of larger changes in added mass on motor learning, we began with the low mass condition for each segment and tested the all-segment loading conditions last. However, it is possible that the chosen order of conditions influenced the study outcomes. Extended exposure to each loading condition could improve our confidence in these findings.

Although knee contact force cannot be measured directly in the native knee, our musculoskeletal modeling approach relied on several assumptions that may limit the accuracy of our contact force estimates. EMG-informed methods have been show to perform well when compared to in vivo measurements from instrumented knee implants [29, 38, 40, 64]. Our model contained 12 major lower-limb muscles, including the major muscles that span the knee joint, but we excluded some smaller muscles (e.g. tensor fasciae latae, sartorius) that contribute to total knee contact force. However, these muscles have small force generating capacities relative to the muscles we chose to include, and collecting surface EMG signals from these muscles can be less reliable.

Furthermore, we used a generic anatomical model that did not incorporate subject-specific skeletal geometry or additional musculotendon parameter calibration [37, 64]. Because our study had a repeated-measures design in which we focused on within-subject changes in contact force, further model individualization may not have had a significant effect on our findings. Finally, we used a simple model of the knee as a single degree of freedom joint. We did not model knee ligaments or the medial and lateral compartments of the knee. Because the majority of individuals with knee OA first experience symptoms in the medial compartment [10], it may be valuable to understanding the frontal plane loading distribution. Future work could increase model complexity by incorporating subject-specific anatomy from medical imaging, modeling the individual compartments of the knee, and calibrating additional musculotendon parameters.

In this study, we estimated changes in peak knee contact force across different lower-limb loading conditions during walking. We used an EMG-informed simulation approach to estimate compressive tibiofemoral contact force from experimental data from 10 healthy adult participants. We found that adding mass to the lower limb increased peak knee contact force, but that the effects were most pronounced for foot loading, which was partially attributed to an increase in stride time. We fit a simple regression to these data to establish relationships between added segment mass and changes in early- and late-stance peak knee contact force. We found that adding mass to multiple segments had a mostly additive effect, but there may be an interaction effect for high mass magnitude for the early-stance peak.

These relationships improve our understanding of the relationship between peak knee loading and added mass on the lower limb. This study also introduces important considerations for individuals with knee osteoarthritis for whom increased knee loading could exacerbate symptoms and disease progression. When evaluating treatment approaches such as footwear, braces, and assistive exoskeletons, we must consider the tradeoff between the expected benefit and adding mass to the lower limbs.

## Conclusion

In this study, we leveraged a combination of experimental data and musculoskeletal simulation to understand the effects of altering lower limb segment masses on peak knee contact force. We collected data from healthy adults walking on a treadmill with varying amounts of mass added to their thighs, shanks, and feet. We found that altering lower limb segment mass has a linear effect on peak knee contact force, but that the slope of this relationship differs across segments and between early- and late-stance.

Adding mass to the feet leads to the greatest increase in peak knee contact force, especially in early-stance. Designers can use these findings to inform the development of wearable devices for individuals with knee osteoarthritis. We invite researchers to use our freely available data and code (https://simtk.org/projects/kcf_leg_mass) to further this work.

## Supporting information

**S1 Fig. EMG waveforms.** EMG waveforms during stance are shown for each loading condition, averaged across participants. EMG data were collected from 9 lower-limb muscles on the right leg. Raw signals were high-pass filtered at 20 Hz, full wave rectified, then low-pass filtered at 6 Hz. Filtered signals were normalized to the maximum seen across maximum voluntary contraction (MVC) and gait trials, then further scaled by during the Calibration stage of EMG-informed simulation. For each muscle, the loaded conditions are shown by color and the unloaded (no mass) condition is shown as a dashed black line.

**S2 Fig. Subject-specific changes in peak knee contact force.** The relationship between added segment mass as a percentage of body weight (% BW) and percent change in peak knee contact force (KCF) is shown for each subject, represented by a different color. Each column indicates a different limb segment loading condition (thigh, shank, foot, all segments). The top row shows changes in early-stance peak KCF, and the bottom row shows changes in late-stance peak KCF.

**S1 Appendix. EMG-informed simulation details and additional results.**

## Acknowledgments

We thank Julie Kolesar for assistance with methodology and software development.

