## Supplemental Appendix 1 for "How peak knee loads are affected by changing the mass of lower-limb body segments during walking"

S1 Appendix

Delaney E. Miller[1*], Ashley E. Brown[1], Nicholas A. Bianco[1], Rucha Bhise[1], Scott L. Delp[1,2,3], Steven H. Collins[1]

**1** Department of Mechanical Engineering, Stanford University, Stanford, California, United States of America
**2** Department of Bioengineering, Stanford University, Stanford, California, United States of America
**3** Department of Orthopaedic Surgery, Stanford University, Stanford, California, United States of America

* dem4[at]stanford.edu

### 1 EMG-informed simulation approach

We created EMG-informed simulations to estimate muscle forces. The optimal control problems were solved using direct collocation in OpenSim Moco [1]. We used the *MocoInverse* interface to formulate the baseline problem with prescribed kinematics. We further customized the problem to add an EMG tracking objective and modify objective weights using the *MocoStudy* interface.

#### 1.1 Modifications to musculoskeletal model

After scaling a generic full-body musculoskeletal model to each subject's static marker data, we made several model modifications to create our EMG-informed simulations. We replaced the 12 Hill-type muscles of the right lower limb in the scaled model with continuously smooth and differentiable muscles [2]. We enabled tendon compliance for all muscles but ignored passive muscle fiber forces, as active muscle forces are the primary contributor to joint moments during walking. Furthermore, uncalibrated passive muscle forces can produce large erroneous contributions to total muscle forces, which could overestimate predictions of knee contact force. We removed the metatarsophalangeal and subtalar degrees of freedom so that these joints remained fixed. For some participants, we did not fix the subtalar joints because an inverse kinematics solution could not be obtained with low marker error, specifically during the swing phase.

We added reserve torque actuators (called "coordinate actuators" in OpenSim) to each joint in the model with an optimal generalized force of 1 (N or Nm) and unbounded control effort. This problem configuration ensured dynamic consistency but discouraged overuse of the reserve actuators at coordinates spanned by muscles. For the knee and ankle flexion coordinates, these actuators provided reserve strength to enforce dynamic consistency despite errors arising from the data. For other joints with no muscles, these actuators provided the full joint moment needed to enforce the dynamic constraints for motion.

#### 1.2 Cost function

The cost function minimized muscle excitation effort and reserve actuator effort while tracking experimental EMG. Each term was integrated over a single stance phase of the right leg with initial time $t_0$ and final time $t_f$. Reserve actuators at the right knee and ankle were heavily penalized, while effort across the remaining reserve actuators was nominally weighted. The cost function can be written as:

$$J = w_1 J_{\text{excitation}} + w_2 J_{\text{knee/ankle}} + w_3 J_{\text{reserve}} + w_4 J_{\text{EMG}} \quad (1)$$

For excitation effort $J_{\text{excitation}}$, we used a *MocoControlGoal* to minimize the sum of squared muscle excitations (12 muscles), integrated over stance:

$$J_{\text{excitation}} = \int_{t_0}^{t_f} \sum_{i}^{12} e_i^2 dt \tag{2}$$

where $e_i$ is the excitation of the $i$th muscle. For knee flexion and ankle flexion reserve actuator effort $J_{\text{knee/ankle}}$, we used a *MocoControlGoal* to minimize the sum of squared coordinate actuator control inputs $u$, integrated over stance. Because we assigned a strength of 1 to each coordinate actuator, each unit of control input corresponded to 1 Newton-meter of torque.

$$J_{\text{knee/ankle}} = \int_{t_0}^{t_f} \sum_{i}^{2} u_i^2 dt \tag{3}$$

The same equation held for reserve actuator effort $J_{\text{reserve}}$ across the remaining 14 pelvis and lower limb coordinates (16 coordinates if the subtalar joints were not fixed). These reserve actuators were instead assigned a small weight $w_3$ to encourage optimizer convergence without contributing substantially to the total cost.

Finally, we implemented a *MocoControlTrackingGoal* to minimize EMG tracking error $J_{\text{EMG}}$. We minimized the sum of squared error between experimental EMG excitation $\hat{e_i}$ and simulated excitation $e_i$ across all 12 muscles and integrated over stance.

$$J_{\text{EMG}} = \int_{t_0}^{t_f} \sum_{i}^{12} (e_i - \hat{e_i})^2 dt \tag{4}$$

The weights on excitation effort and knee and ankle reserve effort varied between calibration and execution (Table S1). During both stages, reserve actuator effort for right knee and ankle flexion $J_{\text{knee/ankle}}$ was penalized such that the muscles were primarily responsible for producing the observed motion. During the calibration stage, higher weights were attached to both excitation effort $J_{\text{excitation}}$ ($w_1 = 10$) and knee and ankle reserve effort $J_{\text{knee/ankle}}$ ($w_2 = 1$). This approach that prioritized excitation effort and reserve effort ensured that co-contraction was minimized and the knee and ankle net joint moments were satisfied almost entirely by muscle forces. Furthermore, the excitation scale factors adjusted the magnitude of the reference EMG signals to most closely produce the desired joint moments.

**Table S1. Cost function weights.**

| Weight | Term | Calibration | Execution |
|---|---|---|---|
| $w_1$ | $J_{\text{excitation}}$ | 10 | 1e-4 |
| $w_2$ | $J_{\text{knee/ankle}}$ | 1 | 0.1 |
| $w_3$ | $J_{\text{reserve}}$* | 1e-4 | 1e-4 |
| $w_4$ | $J_{\text{EMG}}$ | 100 | 100 |

* The hip flexion reserve actuator was assigned a weight of 1e-3.

During execution, the reference EMG signals were scaled by the excitation scale factors found during calibration. The weight on excitation effort $J_{\text{excitation}}$ was reduced to a nominal value ($w_1 =$ 1e-4). The weight on knee and ankle reserve effort was also reduced ($w_2 = 0.1$), which allowed greater deviation from the observed net joint moments and closer tracking of the scaled EMG reference signals. The weight on EMG tracking error $J_{\text{EMG}}$ ($w_4 = 100$) was held constant across Calibration and Execution, since we instead varied the other cost function terms. These weights resulted in a cost function objective for which EMG tracking effort and the remaining terms (excitation and reserve actuator effort) contributed similarly.

For both calibration and execution, a nominal weight (1e-4) was attached to the reserve actuator effort $J_{\text{reserve}}$ for the remaining pelvis and lower limb coordinates. This small penalty yielded the best optimizer convergence while discouraging overuse of the reserve actuators. The right hip flexion reserve actuator was assigned a slightly higher penalty weight (1e-3) due to the biarticular nature of the hamstrings and rectus femoris, although the simulation results were not sensitive to this weight in the region of interest (10-90% of stance).

#### 1.3 Solver settings and computer specifications

We solved each problem in Moco using *MocoCasADiSolver* [3]. Each simulation represented a single stance phase on the right leg from heel strike to toe-off, determined from ground reaction force data. We used a mesh interval of 20 ms. The problem was solved using an optimization constraint tolerance and a convergence tolerance of 1e-4, and the number of iterations was set to a maximum of 5000.

Simulations were conducted on two workstations with Intel i9 processors (24-Core, 5.8 and 6.0 GHz) and 32 GB RAM and were parallelized using Matlab's `parfor` function. Each simulation took approximately 20-30 minutes to solve.

#### 1.4 Experimental EMG tracking error

Experimental EMG signals were tracked closely across conditions and participants. In the region of interest (10-90% of stance), the average RMSE across conditions and participants was 0.021, which represents approximately 2% of the range of possible muscle excitations. EMG tracking RMSE and maximum error by muscle are presented in Table S2. We found that there was higher variation in tracking error across muscles than across participants, with the soleus and vastus lateralis having both the highest RMSE and maximum error. These discrepancies could be explained by a number of causes. Surface electromyography with a single pair of electrodes may not accurately represent the net muscle excitation [4]. The soleus and vastus lateralis have a large volume, which may make a single electrode less reliable. Additionally, we used a generic musculoskeletal model that had several modeling limitations. It is possible that increasing model complexity by incorporating subject-specific geometry [5] or tuning additional musculotendon properties [6–8] would improve EMG tracking performance. Finally, we made several simplifications in our modeling approach, like neglecting smaller leg muscles and passive muscle forces, which could further contribute to a mismatch between experimental EMG and adjusted excitations.

**Table S2. EMG tracking error.** Root mean square error (RMSE) and maximum error between experimental EMG signals and simulation excitations for each muscle between 10 and 90% of stance. The mean and standard deviation across participants is reported for each muscle.

| | **RMSE** | | **Max error** | |
|---|---|---|---|---|
| **Muscle** | **Mean** | **Std dev.** | **Mean** | **Std dev.** |
| soleus | 0.059 | 0.018 | 0.151 | 0.056 |
| gastrocnemius medialis | 0.029 | 0.007 | 0.077 | 0.027 |
| gastrocnemius lateralis | 0.017 | 0.005 | 0.043 | 0.016 |
| tibialis anterior | 0.011 | 0.003 | 0.034 | 0.011 |
| biceps femoris long head | 0.011 | 0.004 | 0.036 | 0.016 |
| vastus lateralis | 0.037 | 0.007 | 0.086 | 0.017 |
| vastus medialis | 0.019 | 0.003 | 0.052 | 0.021 |
| rectus femoris | 0.022 | 0.007 | 0.040 | 0.008 |
| semimembranosus | 0.016 | 0.006 | 0.054 | 0.023 |
| biceps femoris short head* | 0.003 | 0.001 | 0.011 | 0.005 |
| vastus intermedius* | 0.014 | 0.002 | 0.042 | 0.015 |
| semitendinosus* | 0.007 | 0.003 | 0.023 | 0.010 |

*mapped EMG signal

##### 1.5 Knee and ankle reserve actuators

During the region of interest (10-90% of stance), reserve actuators at the knee and ankle produced small amounts of torque. The knee and ankle reserves actuators supplied an average of 5.5e-4 Nm/kg and 4.0e-4 Nm/kg of torque, respectively, normalized to participant body mass. During this same region of interest, the average peak magnitude of reserve torque was 0.0281 Nm/kg at the knee and 0.0089 Nm/kg at the ankle. As a percentage of maximum net joint moment in the region of interest, these values corresponded to 0.06% average and 5% peak at the knee and 0.02% average and 0.5% peak at the ankle. To ensure that timing variation across conditions and participants did not influence the computation of peak values, we first computed the peak value for each participant and each condition, and then averaged across conditions and participants.

Near heel strike and toe-off, we observed larger magnitudes of reserve torque. We attributed these increases to larger errors in our center-of-pressure estimates near heel strike and toe-off. When the vertical component of the ground reaction force is small, center-of-pressure calculations become highly susceptible to measurement error, which can subsequently lead to inaccurate estimates of joint moments. Because the purpose of this study was to evaluate changes in peak knee contact force, which typically occur around 15-40% of stance and 60-90% of stance, we did not explore more complex filtering and thresholding methods to adjust these center-of-pressure calculations near heel strike and toe-off.

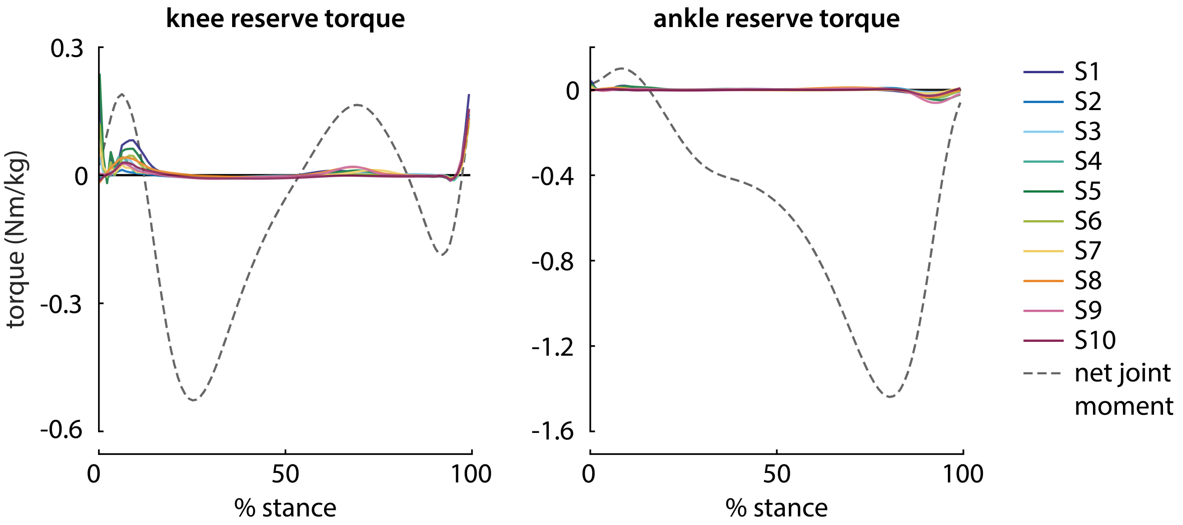


**Fig S3. Knee and ankle reserve torques.** Reserve torques are shown for the stance knee (left plot) and ankle (right plot). Each participant is represented with a different color. Data were averaged across loading conditions.

#### 2 Simulation validation on Grand Challenge subject

We compared knee contact force estimates from our EMG-informed simulation approach to force measurements from an instrumented knee implant. Data were obtained from the Third Grand Challenge Competition to Predict In Vivo Knee Loads [9]. Motion capture, force plate, and electromyography data were synchronized to in vivo measurements of tibial knee load for a single participant walking overground with nine different gait modifications: normal, bouncy, crouch, medial thrust, metatarsophalangeal (MTP) gait, smooth, trunk sway, and two different heights of walking poles. Trials with motion artifact or poor signal quality were excluded, resulting in 32 strides used for analysis.

#### 2.1 Data preparation and simulation

Gait data were prepared for simulation according to the methods described in the main manuscript. Although EMG signals were recorded from 15 muscles on the instrumented limb, we selected signals from 8 muscles to align with the methods used in the main study. The vastus lateralis signal was excluded due to motion artifact; instead, the vastus medialis signal was mapped to the vastus lateralis muscle for simulation. For some strides, EMG signals displaying motion artifact were replaced with an average waveform from other strides of the same gait modification. Each trial was trimmed to one complete stance phase on the instrumented (left) limb. For trials in which the instrumented limb did not strike the middle force plate, a full stance phase was reconstructed using the early-stance phase from the third force plate and the late-stance phase from the first force plate. The first peak of knee contact force was taken between 15 and 40% of the full stance phase, and the second peak between 60 and 90% of stance.

We first performed a Calibration stage to adjust the magnitude of experimental muscle excitations to most closely align with the computed net joint moments. Five strides from normal walking were used for Calibration. Because several MVC signals appeared submaximal, EMG scale factors were allowed to range between 0.33 and 1, rather than between 0.5 and 1 as in the main study. Due to unreliable MVC signals, scale factors for biceps femoris short head and medial gastrocnemius were manually adjusted to better match the scales of adjacent muscles.

Once muscle excitation scale factors were determined, all trials were passed through the Execution stage to estimate muscle forces and, subsequently, total knee contact force. Average RMS EMG tracking error across all muscles and gait conditions was 0.036 in early stance and 0.037 in late stance. Average maximum EMG tracking error across all muscles and gait conditions was 0.044 in early stance and 0.042 in late stance. The average maximum knee reserve torque across all gait conditions was 0.043 Nm/kg in early stance and 0.007 Nm/kg in late stance, or 11.7% and 1.9% of the maximum net joint moment in normal walking, respectively. The maximum ankle reserve torque across all gait conditions was 0.137 Nm/kg in early stance and 0.048 Nm/kg in late stance, or 10.5% and 3.7% of the maximum net joint moment for normal walking, respectively.

#### 2.2 Knee contact force estimates

Simulation estimates of peak knee contact force were comparable to force measurements from the instrumented knee implant for most gait modifications (Fig S4). For normal gait, MTP, smooth, trunk sway, and both short and long walking poles conditions, the average early stance peak KCF error was 0.29 $\pm$ 0.23 BW. In the bouncy, crouch, and medial thrust conditions, our simulation approach overestimated the early-stance peak of knee contact force. For these conditions, average early stance peak KCF error was 1.62 $\pm$ 0.49 BW. The average late-stance peak KCF error across all conditions was 0.14 $\pm$ 0.24 BW. Crouch gait was excluded from the late-stance KCF error calculation because it lacked a distinct late-stance KCF peak in both the measured force implant data and simulation-estimated contact force.

It is unclear why our simulation overestimated early-stance peak contact force for some gait conditions. In these conditions, we observed large experimental muscle excitations for the vastus medialis in early stance, which were consistent with the large peak knee extension moments calculated through inverse dynamics. It is possible that the overestimates of peak knee contact force arise from errors in the anatomical model, such as inaccuracies in muscle attachment points or the location of the knee joint center. The largest errors between simulation estimates and implant measurements of peak knee contact force occurred in conditions with more knee flexion in early stance. These differences may have arisen due to modeling inaccuracies that became more apparent with more extreme knee kinematics.

#### 2.3 Pairwise ranking accuracy

In addition to evaluating the absolute errors in peak knee contact force, we evaluated how well the EMG-informed simulation approach ranked gait modifications by peak knee contact force. For every possible pair of gait modifications, for early- and late-stance separately, we compared the simulation estimated peaks in knee contact force to the measured peaks from the instrumented knee implant. If simulation correctly

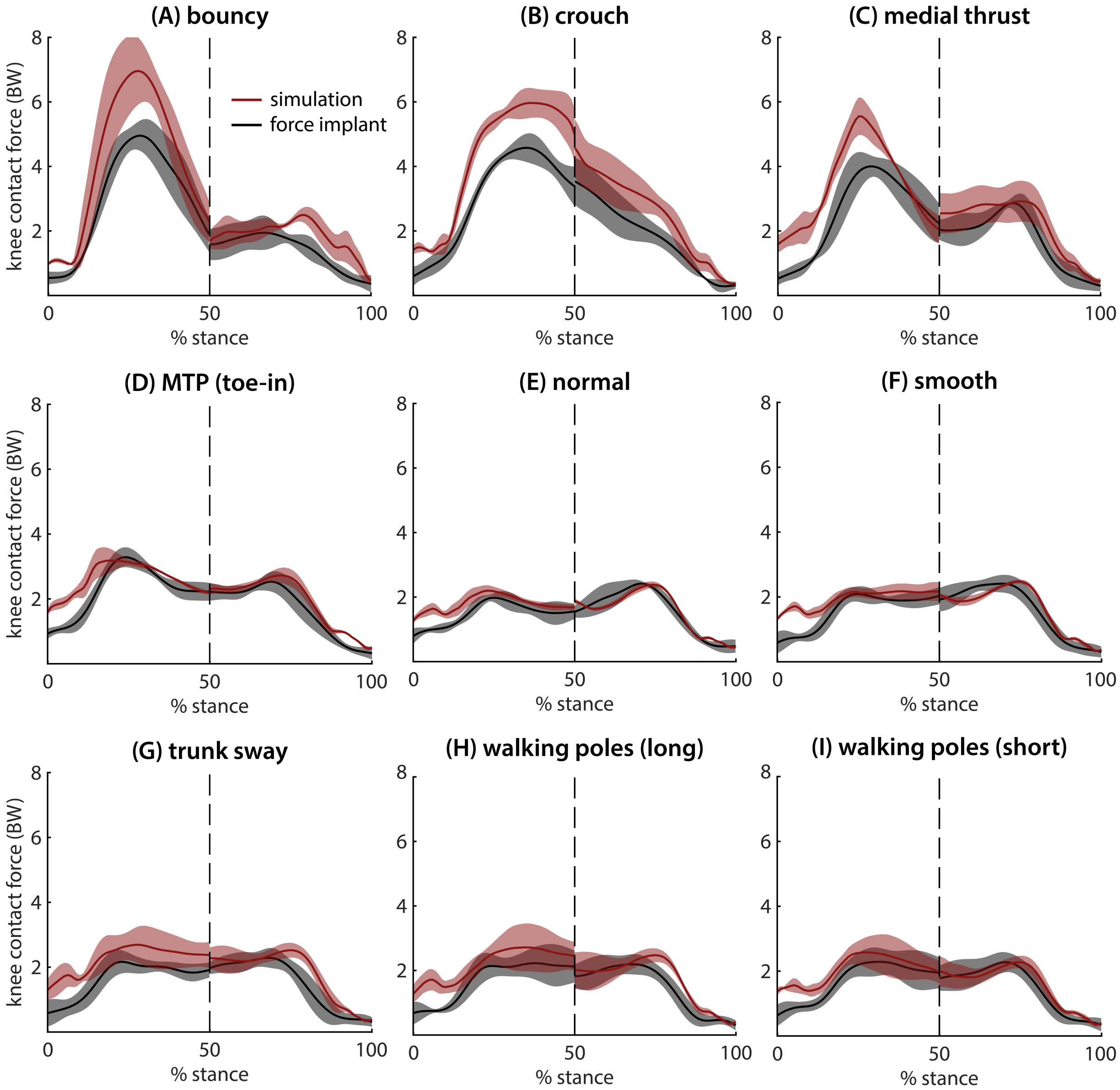


**Fig S4. Simulation-estimated versus measured knee contact forces.** Average waveforms during stance of measured knee contact force from instrumented knee implant (black) and estimated knee contact force from EMG-informed simulations (red) are shown for each gait modification: (A) bouncy, (B) crouch, (C) medial thrust, (D) MTP (toe-in), (E) normal, (F) smooth, (G) trunk sway, (H) long walking poles, and (I) short walking poles. Standard deviation across strides is shown in the shaded regions.

identified the larger peak in knee contact force, the comparison was assigned an accuracy of 1, and 0 otherwise. We excluded comparisons for which the true difference between KCF peaks was less than 0.1 BW, as these small differences may not be clinically meaningful. The EMG-informed simulation estimates achieved pairwise ranking accuracy of 92% for early-stance peak contact force and 81% for late-stance peak contact force. This result indicates that our simulation approach correctly identified directional changes in peak knee contact force when compared to instrumented knee implant data.
