## Supplementary figures and images for "How peak knee loads are affected by changing the mass of lower-limb body segments during walking"

### Supplemental Figure 1

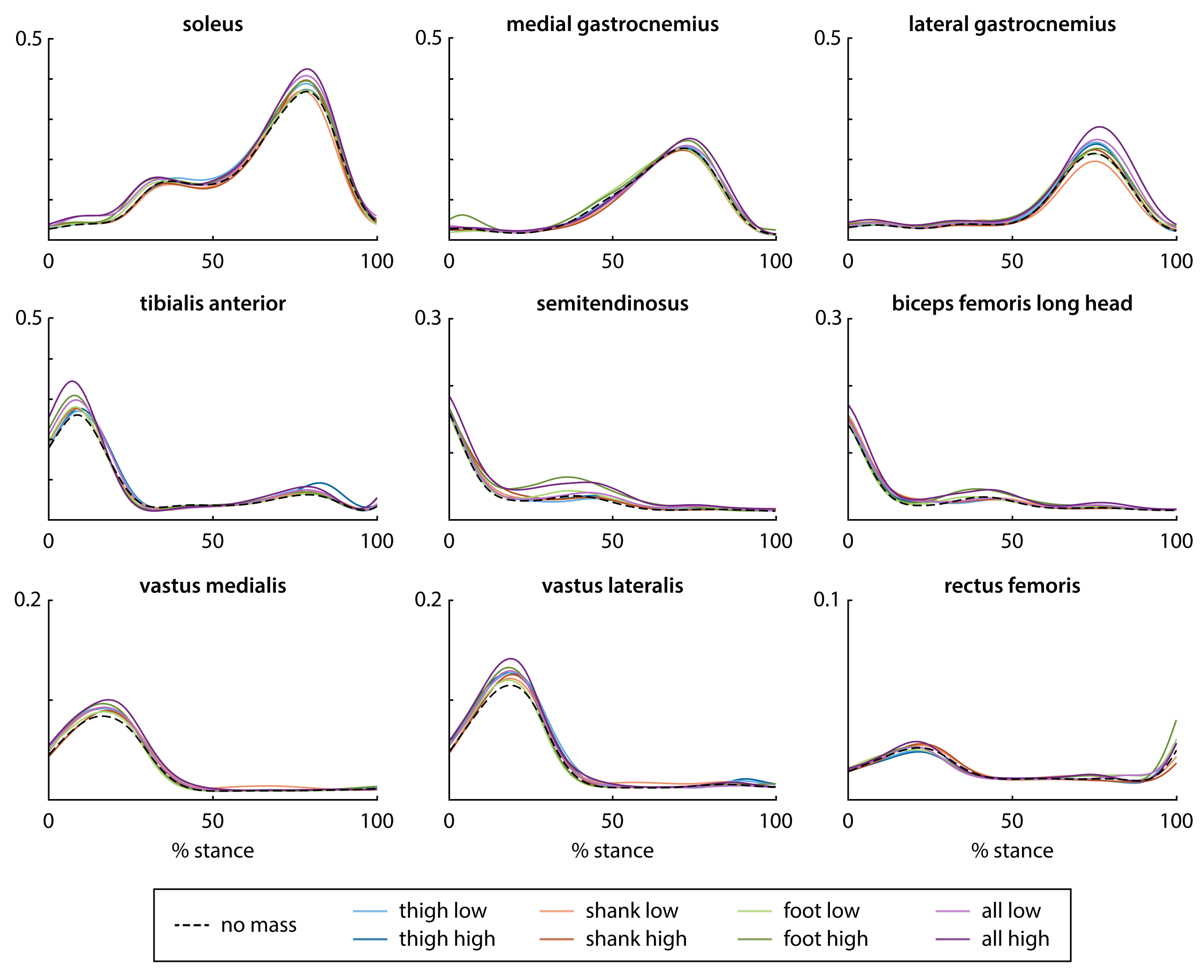

### Supplemental Figure 2

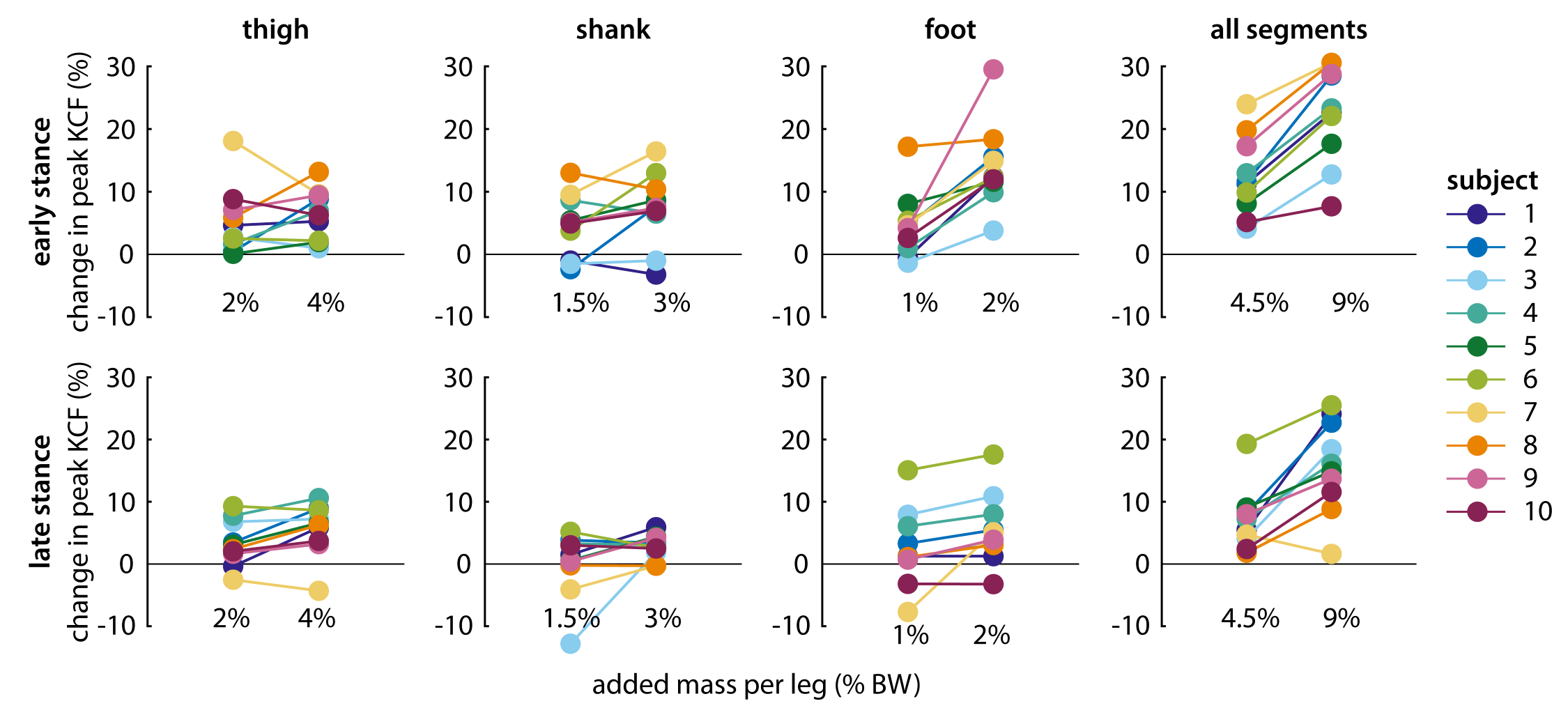
